# Factors underlying creativity show distinct genetic architecture and genetic correlations with psychiatric disorders

**DOI:** 10.64898/2026.09.30.755774

**Authors:** Penghao Xia, Laura W. Wesseldijk, Giacomo Bignardi, Yi Lu, Fredrik Ullén, Miriam A. Mosing

## Abstract

While creativity is widely recognized as multifaceted, genome-wide association studies have largely focused on individual measures. Here, we used genomic structural equation modeling of six creativity-related GWASs and identified two latent genetic factors: a general factor and an artistic factor, with 22 and 13 genome-wide significant loci, respectively, nine of which are novel. The general factor was almost perfectly genetically correlated with educational attainment (*r*_g_ = 0.96), suggesting that it primarily captured genetic influences shared with cognitive and educational processes rather than creativity-specific variation. In contrast, the artistic factor showed only a weak genetic correlation with educational attainment (*r*_g_ = 0.11) and consistent positive genetic correlations with psychiatric disorders, whereas associations for the general factor were largely null or negative. Biological annotation analyses further identified distinct enrichment patterns unique to the artistic factor, providing initial evidence that different dimensions of creativity may be characterized by partially distinct neurobiological pathways.

## Introduction

Creativity has long been recognized as a driving force behind cultural and societal processes^1^, yet our understanding of this human ability remains limited. Numerous measures of creativity have been developed, including the Creative Achievement Questionnaire^2^ and the Divergent Association Task^3^, but there is little consensus on how creativity should be best assessed^4^. The diversity of measures raises the question whether they capture a common underlying construct or distinct facets of creativity. Consistent with the latter possibility, previous research suggested that inconsistent associations between creativity and mental health may partly reflect heterogeneity in creativity measurement^5–9^.

Since creativity is a heritable trait (with twin-study-based heritability up to 70% across measures^10–12^), researchers started to explore the connection between creativity and mental health from the genetic perspective. Indeed, an earlier effort^13^ exploring the genetic link between creativity and mental health using genotype data, showed that creative individuals had higher polygenic scores (PGSs) of schizophrenia and bipolar disorders. Lately, researchers started to conduct large-scale genome-wide association studies (GWASs) to identify the genetic architecture of creativity and related traits. These studies identified some common single-nucleotide polymorphisms or SNPs related to creativity (SNP-based heritability ranging from 0.07 to 0.22), and revealed substantial genetic overlap with psychiatric disorders (e.g., genetic correlations approaching 0.4)^14–17^. However, findings varied largely depending on how creativity was defined and measured; for example, significant associations between creativity and schizophrenia were found in some GWASs^14,17^ but not others^16^. Psychiatric disorders are differentiated into distinct dimensions whose genetic overlap and specificity have been thoroughly investigated in recent years^18–20^. In contrast, previous GWASs of creativity have treated creative outcomes either as a monolithic category or in isolation. This is at odds with the idea that creativity is a multifaceted construct^4,21^, and may explain the inconsistent phenotypic and genetic overlap between creativity and other traits reported in previous studies.

One solution is to examine the genetic overlap across diverse measures, rather than just focusing on a single one. A recent twin study^21^ suggested distinct genetic factors underlying various creativity measures: a general genetic factor underlying all creativity measures - artistic (i.e., artistic achievement and three openness subscales) and non-artistic (i.e., scientific achievement, occupational creativity, and divergent thinking), and a second genetic factor shared only among artistic measures. Informed by the findings of this twin study, we used *genomic structural equation modeling* (Genomic SEM)^22^ to validate the two-factor model with a preregistered (https://aspredicted.org/k9ph8a.pdf) selection of GWASs on creativity-related measures: occupational creativity (*N* = 241,736)^16^, artistic (*N* = 64,580) and scientific occupations (*N* = 76,180)^14^, education in arts (and humanities; *N* = 463,134) and natural sciences (with mathematics and statistics; *N* = 463,134)^15^, as well as openness (*N* = 504,063)^17^. We then conducted a multivariate GWAS on each of the two latent creativity factors by synthesizing univariate GWASs on the single creativity-related measures. To validate the results, we derived PGSs for the latent factors and examined their performance in predicting various creativity measures in an independent Swedish sample^23–26^. We compared the two latent factors in terms of their genetic correlations with psychiatric traits, as well as intelligence and educational attainment. Finally, to provide initial biological grounding for these associations, we examined the enrichment profiles of the genes associated with the two factors.

## Results

### Genomic SEM confirmed two latent factors

We fit the suggested two-factors solution model and ran multivariate GWASs by incorporating the summary statistics from GWASs on occupational creativity^16^, openness^17^, artistic occupation^14^, education in arts^15^, scientific occupation^14^, and education in natural sciences^15^. A summary (including sample size and heritability) of the univariate GWASs (i.e., GWASs on single creativity-related measures) is in Supplementary Table 1. Genetic correlations (*r*_g_) from linkage disequilibrium (LD) score regression (LDSC)^27^ revealed genetic overlap across all creativity measures (*r*_g_ values ranging from 0.38 to 0.92; Fig. 1a).

**Fig. 1.**
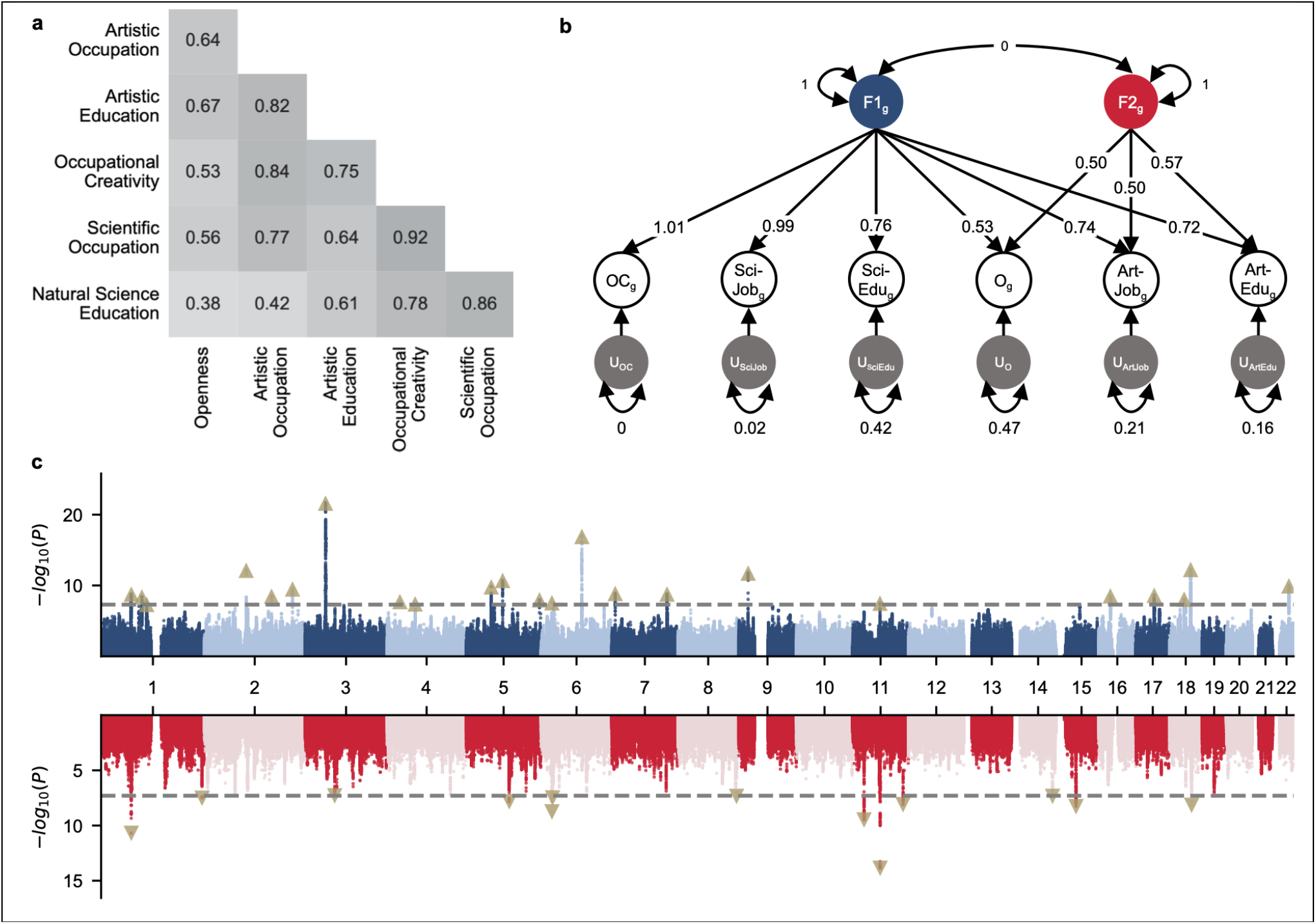
Shared genetic structure across creativity-related measures. **a**, genetic correlations between the six creativity-related measures. All genetic correlations were significant after FDR correction. **b**, genome-wide structural equation model. The model extracts a general factor (F1_g_) and an artistic factor (F2_g_) from the genetic influences underlying the creativity-related measures of occupational creativity (OC_g_); scientific occupation (Sci-Job_g_); education in natural sciences, mathematics, and statistics (Sci-Edu_g_); openness (O_g_); artistic occupation (Art-Job_g_); and education in arts and humanities (Art-Edu_g_). The reported estimates of the factor loadings (single-headed arrows) and residual variances of the creativity-related measures (double-headed arrows on the circles labeled with “u”) were standardized relative to SNP-based heritability in the model. The model fixes the residual variance of occupational creativity as 0, the variances of latent variables (double-headed arrows on F1_g_ and F2_g_) to be 1 and their covariance (the double-headed arrow between factors) to be 0. **c**, Miami plot of the two multivariate GWASs on the general and artistic factors. The plot shows -*log*_10_*P* of each SNP’s association with the general and the artistic factor. The dashed lines represent the genome-wide significance threshold of *P* = 5×10^−8^. Significant genomic loci are visualized as triangles.

To confirm the presence of two genetic factors explaining the genetic overlap across creativity measures^21^, we ran genomic Confirmatory Factor Analysis (CFA) with Genomic SEM. The genomic CFA indicated a good fit (SRMR = 0.0500, CFI = 0.9849) of the two-factor structure (details of parameter estimates in Supplementary Table 3)^28^: Factor 1, for simplicity hereafter referred to as the *general factor*, underlying all creativity-related measures and Factor 2, the *artistic factor*, shared only by artistic creativity measures. In the initial modeling, occupational creativity had a negative but non-significant standardized residual variance of 0.04 (*P* = 0.26, see Supplementary Table 3). Therefore, we then fit the two-factor model with the residual variance of occupational creativity constrained as 0 (Fig. 1b; details of parameter estimate in Supplementary Table 3). As the constrained model showed a similar model fit (SRMR = 0.0502, CFI = 0.9850) as the initial model, we based all the follow-up analyses on the constrained model. To obtain SNP-based associations with each of the two identified factors, we used Genomic SEM to run two multivariate GWASs, based on the 2-factor model verified by the genomic CFA (Fig. 1c). We identified 23 and 13 genomic loci (locus definition in Methods) for the general and artistic factors, respectively (Supplementary Tables 4 and 5), of which nine were novel or not discovered in the creativity GWASs included in the Genomic SEM (six for the general factor and three for the artistic factor).

### PGS validation

To validate the multivariate GWAS results, we examined if the PGSs of the two factors could predict a range of creativity phenotypes in the STAGE cohort^23–26^ (sample size ranged from 3,379 to 4,662 across nine creativity measures including creative achievements, divergent thinking, and openness subscales), a sample independent of the discovery univariate GWASs. As illustrated in Fig. 2 (details in Supplementary Table 6), the PGS of the general factor (underlying all creativity-related measures in the Genomic SEM) showed pervasive positive associations with nearly all creativity measures, except for the openness subscale of creative imagination and artistically creative occupation. In comparison, a higher PGS of the artistic factor was mainly associated with higher scores in artistic creativity measures (i.e., the openness subscales and artistic achievement, with the exception of artistically creative occupation).

**Fig. 2.**
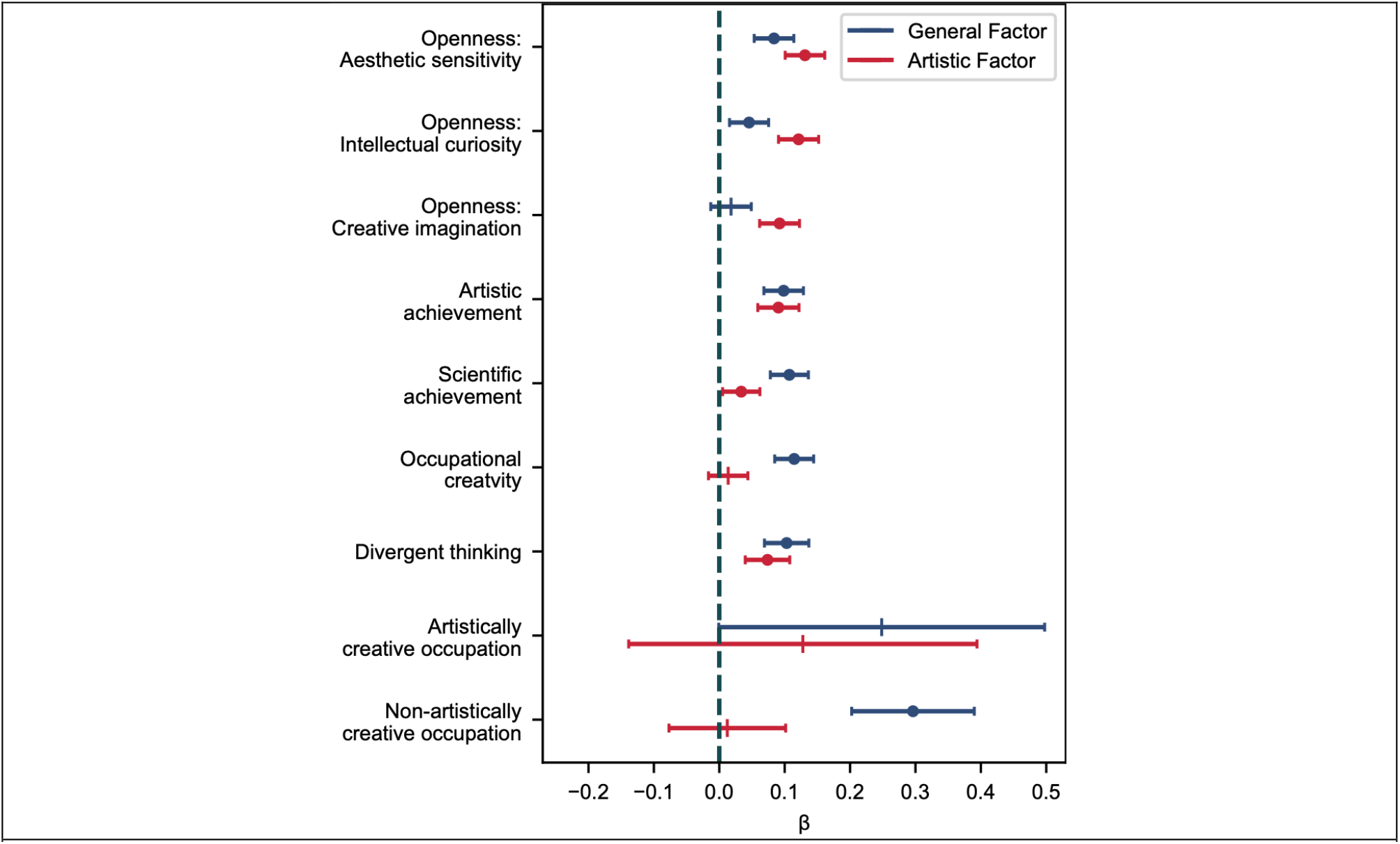
Predictability of creativity factor polygenic scores (PGSs). The plot illustrates the regression coefficients (β and 95% confidence intervals) of PGSs of the general factor and the artistic factor predicting continuous (standardized) and binary creativity measures (artistically and non-artistically creative occupations) in the STAGE cohort. Point estimates that are significant after false-discovery rate correction for 18 tests are plotted as dots.

### Genetic correlations with psychiatric disorders, intelligence, and education

We used LD score regressions^27,29^ to calculate the genetic correlations of the general and artistic factors with psychiatric disorders. For comparison, we also calculated genetic correlations between the individual creativity-related measures and psychiatric disorders. Additionally, we examined the genetic correlations of the two factors with intelligence and education attainment. Supplementary Table 7 lists the GWASs on psychiatric disorders, intelligence, and education attainment.

As shown in Fig. 3 (details in Supplementary Table 8), higher scores on the artistic factor were associated with higher genetic risks of all psychiatric disorders (with *r*_g_ ranging from 0.23 to 0.38), including genetic correlations that were not detected with single creativity-related measures, such as with attention-deficit/hyperactivity disorder (ADHD; *r*_g_ = 0.30, *s*.*e*. = 0.03) and substance use (*r*_g_ = 0.25, *s*.*e*. = 0.04). In contrast, the general factor had mainly non-significant or negative genetic correlations with psychiatric disorders, with significantly negative genetic correlations ranging from -0.53 to -0.19. The only exceptions were weakly positive genetic correlations with bipolar disorder (*r*_g_ = 0.10, *s*.*e*. = 0.03) and the compulsive factor (underlying obsessive-compulsive disorder, anorexia nervosa, and Tourette’s syndrome^30^; *r*_g_ = 0.09, *s*.*e*. = 0.04), and a moderately positive genetic correlation with autism (*r*_g_ = 0.21, *s*.*e*. = 0.04).

**Fig. 3.**
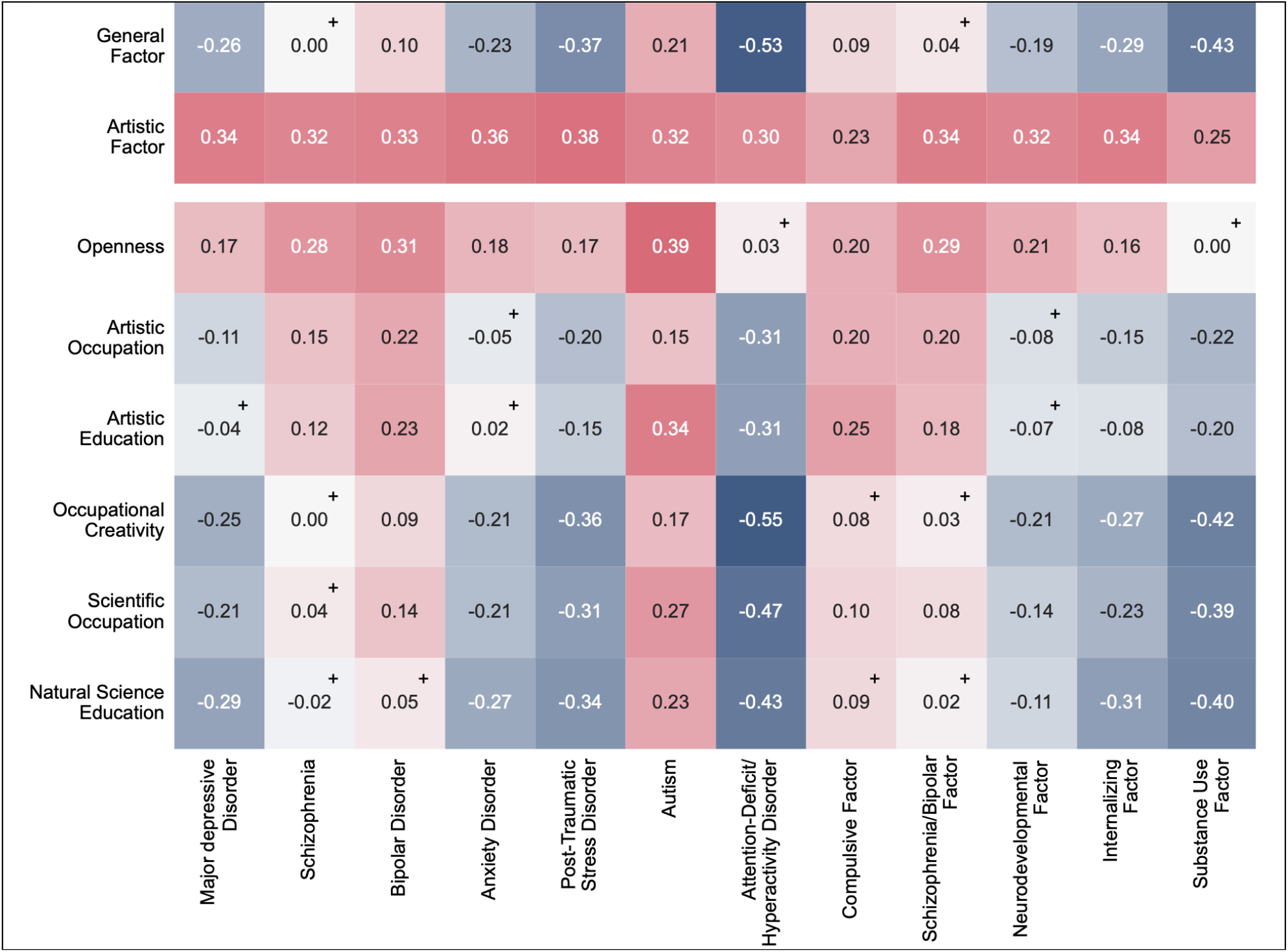
Genetic correlations between creativity and mental health phenotypes. This plot visualizes the genetic correlations (i.e., values in the cells) of the general factor, the artistic factor, and creativity-related measures with mental health phenotypes. Positive and negative genetic correlations are represented in red and blue, respectively. Non-significant genetic correlations (after FDR correction) are marked with “+”.

As for the non-psychiatry phenotypes (see Supplementary Table 8), while the artistic factor only had a low genetic correlation (*r*_g_ = 0.11, *s*.*e*. = 0.02) with educational attainment (i.e., number of education years) and a non-significant genetic correlation (*r*_g_ = 0.07, *s*.*e*. = 0.03) with intelligence, the general factor showed strong genetic correlations with educational attainment (*r*_g_ = 0.96, *s*.*e*. = 0.01) and intelligence (*r*_g_ = 0.74, *s*.*e*. = 0.02). Despite the high genetic correlation, the general factor had two significant genomic loci (with the top SNPs being rs61792940 and rs6519190; details in Supplementary Table 4) that were not in the educational attainment GWAS.

### Biological annotation

To further explore the biological mechanisms underlying the latent factors, we conducted gene-based, gene-set, and enrichment analyses using MAGMA^31^ integrated in FUMA^32^. As a default setting of FUMA, Bonferroni correction was applied to each creativity factor to account for multiple testing in each analysis. Our gene-based analyses (Supplementary Table 9) identified 103 (e.g., DCC, MST1, MST1R, TCTA, and AMT) and 35 genes (e.g., BDNF, LRFN4, SEMA6D, NRXN2, SYT12, FOXP1, and CLOCK) significantly associated with the general and the artistic factor, respectively. Across the 17,005 tested gene-sets, effective genes for the general factor were enriched in three gene-sets affecting synapses (significant gene-sets in Supplementary 10) but no gene-set was significant for the artistic factor.

Among the 54 tissues in the Genotype-Tissue Expression (GTEx v8) dataset^33^, effective genes were significantly enriched in only brain tissues for both factors (Fig. 4a; complete results in Supplementary Table 11). Especially, genes associated with the artistic factor were enriched in the spinal cord and the substantia nigra, but the genes associated with the general factor did not. To further explore gene expression in the nervous system, we used single-cell gene expression profiles in different brain regions and spinal cord from three postmortem adults^34^ to test if genes with higher expression in one cell-type are more strongly associated with the factors underlying creativity measures (when the average gene expression level across cell-types in the same region is controlled for), at the alpha level of 0.05/1,135 (i.e., the number of combinations of regions and cell-types). As shown in Fig. 4b (showing only a subset of cerebral cortex regions and cell types for visual clarity) and Supplementary Table 12 (with all regions and cell types), associated genes were more enriched in neurons than other cells for both factors. Additionally, genes associated with the artistic factor were also more enriched in oligodendrocyte precursor cells.

**Fig. 4.**
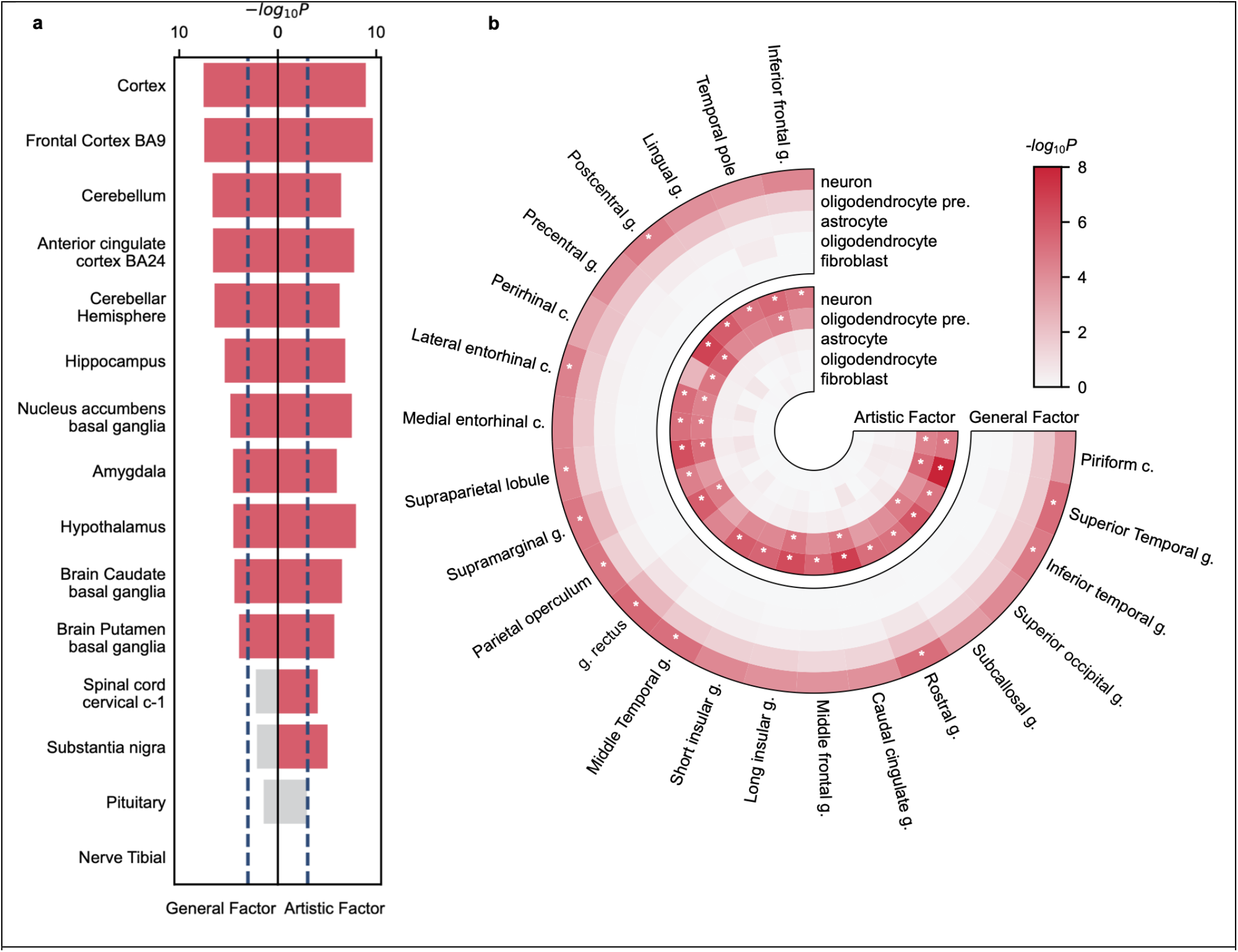
Enrichment of genes associated with factors underlying creativity. This figure illustrates the -*log*_10_*P* values of a selected set of tissues and cell-types in the nervous system, with complete results in Supplementary Tables 11 and 12. **a**, MAGMA tissue expression analysis results. Dashed lines indicate the *P*-value threshold after Bonferroni correction for the 54 GTEx v8 tissues (with *P* ≤ 0.05/54). Significant results are plotted as red bars. **b**, MAGMA cell-type expression analysis results. Results that were significant after Bonferroni corrections (with *P* ≤ 0.05/1,135) are marked with asterisks. Oligodendrocyte pre., oligodendrocyte precursor cell; g., gyrus/gyri; c., cortex.

## Discussion

By synthesizing the six GWASs on creativity-related measures with genomic SEM, this study confirmed two latent genetic factors underlying creativity and uncovered their genetic architecture. The first, a general creativity factor, was shared between all measures, providing evidence for shared genetic influences across diverse indicators of creativity despite their heterogeneity. In addition, a second factor was specific to measures reflecting artistic creativity, indicating genetic markers only relevant to the arts. Notably, the artistic factor showed consistently positive genetic correlations with several psychiatric disorders, including schizophrenia, autism, and substance use. In contrast, genetic correlations between the general creativity factor and psychiatric disorders were mostly negative or non-significant. Furthermore, the general factor, but not the artistic factor, largely overlapped with genetic influences on educational attainment, suggesting that much of its genetic variance reflects influences shared with educational and cognitive outcomes. Finally, tissue and cell-type enrichment analyses revealed distinct patterns of biological enrichment for the artistic factor, pointing to potentially different neurobiological mechanisms underlying artistic creativity and its positive genetic associations with psychiatric disorders.

### Confirming two creativity factors differentially related to psychopathology and cognition

In line with the previous twin study^21^, genomic SEM showed a good fit of a two-factor model to explain the shared genetic architecture underlying different creativity indicators: a general creativity factor and a factor specific to artistic creativity. Interestingly, the multivariate GWASs demonstrated that these two factors were distinctively related to mental health and education.

The artistic factor showed consistent positive genetic correlations with psychiatric disorders. Whereas the general factor exhibited mostly negative or non-significant genetic correlations with psychiatric disorders, with the exceptions of bipolar disorder, autism, and the compulsive disorder factor. In contrast, the general factor was strongly genetically correlated with intelligence and especially educational attainment, while the artistic factor showed little genetic overlap with either trait. In line with the general factor’s genetic correlations with intelligence and educational attainment, the GWASs of occupational creativity^16^ and creative occupations^14^ also reported substantial genetic correlations with intelligence (0.46 - 0.72), and near unity correlations with educational attainment (0.74 - 0.93). Since intelligence^38^ and educational attainment^39,40^ are themselves genetically associated with lower risk of several mental disorders, including major depressive disorder and ADHD, these traits may partly explain the negative genetic correlation between the general factor and psychiatric disorders. Autism appears to be an exception to this pattern: the positive genetic correlation between the general factor and autism instead reflect the well-established positive genetic associations between autism and intelligence^38^, as well as cognitive characteristics linked to both traits, such as heightened attention to detail^41^.

The two factors could offer a potential explanation for the heterogeneous associations between creativity and mental health reported in previous research^5–9^. In line with this, positive associations between creativity and psychopathology have been most frequently reported by studies utilizing indicators of artistic creativity, such as artistic occupations^13,42^, education^43^, or achievements^5,44^. For example, Kyaga and colleagues^42,45^ found unipolar depression to be overrepresented among individuals with artistic occupations (a measure loaded on both the artistic and general factors), but underrepresented among those with scientific occupations (a measure loaded only on the general factor).

Notably, ADHD and the substance use factor had positive genetic correlations with the artistic factor despite showing no or negatively significant genetic correlations with any individual creativity-related measures. These associations could only be revealed after modeling the shared genetic architecture among multiple creativity indicators, highlighting the value of multivariate approaches and the importance of incorporating diverse creativity measures when investigating its genetic basis.

### So, what do the genetic factors represent?

For the general factor, one possibility is that it reflects the educational and cognitive demands inherent in creativity-related measures based on occupations and educational outcomes. In other words, genetic variants related to higher educational attainment are likely more important in some creative domains (e.g., science) compared to others, and those genetic variants may partly counteract the higher psychiatric risk shared with genetic variants related to higher artistic creativity. Under this interpretation, future GWASs employing more objective measures of creativity, such as divergent thinking or creative achievement, may reveal a factor less strongly tied to educational attainment. Supporting this interpretation, phenotypic studies measuring creativity with creative achievements^11,46^ and divergent thinking tests^47,48^ reported phenotypic correlations (0.17 to 0.44) with intelligence that were smaller than the genetic correlations between occupation-based creativity and intelligence (between 0.46 and 0.72).

Alternatively, the general factor may represent a genuine component of creativity that is inherently associated with cognitive ability. The long-standing debate regarding the relationship between intelligence and creativity is consistent with this interpretation, with some theories proposing that cognitive ability is a prerequisite for creative performance^48^, or even essentially creativity^49^. Importantly, however, our multivariate GWAS identified two genomic loci for the general factor that did not overlap with loci identified in the large GWAS of educational attainment^39^, suggesting that the factor cannot be reduced entirely to educational attainment.

The artistic factor, in contrast, was extracted from artistic creativity measures after accounting for variance shared across all creativity measures. Because it showed little genetic overlap with intelligence or educational attainment, it may capture characteristics that direct creative expression specifically toward artistic domains, such as artistic interests or sensitivity.

Alternatively, it may capture psychological traits linked to increased psychiatric vulnerability, given its consistently positive genetic correlations with a broad range of psychiatric disorders.

Regardless, future GWASs on creativity should incorporate more creativity(-related) measures (e.g., divergent thinking) apart from occupational and educational proxies, despite the inconvenience in data collection from large samples.

### Neurobiological links between genes and creativity

Genes associated with both the general and the artistic factors were significantly enriched exclusively in brain tissue (Fig. 4a); no non-neural tissue reached significance after correction. Notably, within the nervous system, both factors showed preferential enrichment in neurons relative to other cell types, (Fig. 4b and Supplementary Table 12). This pattern appears unsurprising for a behavioral trait but supports that creativity-related genetic variation acts primarily through neural mechanisms involved in creative cognition and achievement, as would be expected.

More interestingly, genes associated with the artistic factor, but not the general factor, were enriched in the substantia nigra. The nigrostriatal dopaminergic pathway has been linked to creative cognition, and more specifically, regulation of cognitive flexibility, i.e., divergent thinking, broad attentional scope, and a flexible switching between different perspectives^50^. Furthermore, dopaminergic dysregulation within this system is a key feature of psychiatric conditions that genetically correlate with the artistic factor, in particular schizophrenia^51^. The selective enrichment in substantia nigra, therefore, suggests that functional properties of this brain region may be one neurobiological link between artistic creativity and psychiatric liability.

Moreover, cell-type enrichment analyses further supported the distinction between the two factors. Compared to genes associated with the general factor, those underlying the artistic factor were additionally enriched in oligodendrocyte precursor cells, whose canonical role is to differentiate into myelinating oligodendrocytes^52^, which form myelin sheaths around neuronal axons in the white matter^53,54^). Myelination by oligodendrocytes in the brain is moderated by experience and neuronal activity, and such myelin plasticity has been proposed to be involved in learning^55,56^. In line with the role of white matter plasticity in cognition, a few brain imaging studies have associated creativity with white matter integrity (i.e., axonal coherence and myelination)^57,58^. At the same time, oligodendrocyte lineage cells and myelination have been associated with multiple psychiatric disorders, including major depressive disorder^59^, schizophrenia^60^, and post-traumatic stress-related disorder^61^, and autism^62^. Although specific mechanisms would remain highly speculative, the present findings suggest the possibility that neurobiological links between genes and creativity also may involve oligodendrocyte glia cells.

Finally, we note that the specific genes significantly associated with artistic creativity include the gene for the brain-derived neurotrophic factor (BDNF). The central roles of this neuropeptide in neuronal plasticity and synaptic development have been intensively studied for many years in regions like frontal, hippocampal, and amygdala networks^36,37^. More specifically, BDNF signaling is also important for homeostatic plasticity, i.e., plastic mechanisms that regulate neural activity around a target set-point; dysfunctional homeostatic plasticity has in turn been proposed as a mechanism of psychiatric disease^63^, such as schizophrenia and bipolar disorder^35^.

### Limitations

Finally, several limitations are inherent to the GWAS and Genomic SEM framework.. Because our analyses relied on common SNPs, they capture only a portion of the heritable variation underlying creativity-related traits and do not account for other sources of genetic influence, such as rare variants, structural variants, or non-additive genetic effects. In addition, Genomic SEM models the genetic variance shared across measures and therefore does not capture measure-specific genetic influences, nor does it provide information on environmental sources of variation. As a result, the factors identified here should be interpreted as reflecting shared common-SNP genetic architecture rather than the complete etiology of creativity. Furthermore, because the underlying GWASs were conducted primarily in individuals of European ancestry, the generalizability of these findings to other populations remains to be established. Next, because several of the GWASs in the Genomic SEM assessed occupational and educational indicators of creativity, the general factor may largely reflect genetic influences shared with educational attainment. Future GWASs on more objective measures of creativity (e.g., divergent thinking) are needed to better understand creativity’s genetic architecture. Second, when identifying unique or novel genomic loci, we compared our multivariate GWAS results to univariate GWASs on openness and educational attainment with 23andMe samples excluded.

Because differences in sample composition and statistical power can influence locus discovery, some loci classified as unique here may reflect differences between the available GWAS datasets rather than creativity-specific genetic effects.

## Conclusion

Based on the existing GWASs on creativity-related phenotypes, the current study identified two distinct latent genetic factors underlying creativity that were differentially related to cognition and psychopathology. Particularly, the artistic, but not the general, factor showed consistently positive genetic correlations with various psychiatric disorders, whereas the general factor exhibited an extremely high genetic correlation with educational attainment. These distinct genetic profiles may explain why previous efforts to characterize creativity and its associations with other traits have yielded inconsistent results. Finally, tissue and cell-type enrichment analyses highlighted brain regions and cell-types that may play a role in artistic creativity and its genetic overlap with psychiatric disorders, providing targets for future validation in neuroimaging and related studies. Together, these findings suggest that creativity is genetically multidimensional and that different facets of creativity may arise from partly distinct biological pathways.

## Methods

The study and analyses were approved by the Swedish Ethical Review Authority (Dnr 2020-02575, 2021-02014, 2022-00109-02, 2023.04985.02).

### Preparation of summary statistics

A quality control procedure was applied to all GWAS summary statistics before running Genomic SEM and genetic correlations. Specifically, we dropped genetic variants on sex chromosomes, being insertion-deletion, without reference SNP cluster identification (rsID), or with minor allele frequencies (MAF) < 0.01. When the relevant information was provided, we excluded SNPs with imputation quality < 0.7 or with significant heterogeneity (i.e., *P* of Q-statistics ≤ 5e-8).

Apart from quality control, each SNP was harmonized to have the same effective allele (and non-effective allele) across all summary statistics. Using the criterion in the R package TwoSampleMR^64^, we removed palindromic SNPs with MAF > 0.42 due to ambiguity. Following the manual of Genomic SEM, we calculated the effective sample sizes (when not available) for GWASs on binary variables^65^ or from Genomic SEM^66^. Additionally, for creativity GWASs entering Genomic SEM, we selected the common SNPs across the studies.

### Genomic SEM

#### Genome-wide models

To validate the latent factor structure of creativity from a previous study^21^, we used Genomic SEM and build a two-factor genome-wide model (more specifically, a genomic CFA, genomic Confirmatory Factor Analysis) on the summary statistics of GWASs on creativity-related measures: occupational creativity^16^ and openness^17^, artistic occupation^14^ and education in arts^15^ (corresponding to artistic achievement and aesthetic interest in the previous study^21^), as well as scientific occupation^14^ and education in natural sciences^15^ (corresponding to scientific achievement in the previous study^21^). Following the suggested criteria^28^, a model fits good if its SRMR (Root Mean Square Error of Approximation) ≤ 0.05 and CFI (Comparative Fit Index) ≥ 0.97.

As the publicly available openness GWAS^17^ incorporated the STAGE cohort, which served as the validation sample in the current study, we requested and used GWAS for openness, leaving out the STAGE cohort (23andMe was also excluded due to access restriction). The summary statistics were first munged by LDSC^29^ and restricted to SNPs in HapMap3^67^. We then used the R package “GenomicSEM” to fit the genome-wide model in Fig. 1b. Since Genomic SEM requires population prevalences for binary measures, we used the prevalences reported in the respective GWASs (see Supplementary Table 1).

#### Multivariate GWASs

After validating the two-factor model, we further ran multivariate GWASs (which synthesize univariate GWASs on individual creativity-related measures) with Genomic SEM to estimate the effects of SNPs on the two latent factors. The summary statistics of the multivariate GWASs then underwent the same quality control procedure as the GWASs on the separate creativity-related measures. Additionally, we excluded SNPs for which the model might not have converged (as suggested by the warning messages), and whose Z scores changed more than 0.025 after smoothing. Effective sample sizes were derived based on SNPs with MAF ≥ 0.1^30,68^.

#### Locus definition

Significant loci were defined by FUMA^32^ with the LD structure pre-calculated in the European sample from the 1000 Genome Project Phase 3^69^. Among the genome-wide significant SNPs (*P* ≤ 5e-8), less correlated SNPs (*r*^2^ < 0.6) were regarded as independent. Two LD blocks of independently significant SNPs were merged into a genomic locus if their distance was smaller than 250 kb. Before defining genomic loci, we excluded non-significant SNPs (*P* > 0.05 and MAF < 0.01), or on the MHC (major histocompatibility complex) region. To identify novel genomic loci, we applied the same locus definition method to the univariate GWASs on creativity-related phenotypes after SNP quality control. When defining the genomic loci associated with openness, we used the GWAS with STAGE (but without 23andMe), rather than the one (without STAGE or 23andMe) used in Genomic SEM. A significant genomic locus in our multivariate GWASs was regarded as novel when it did not overlap with any significant loci in the univariate GWASs. We used the same strategy to identify genomic loci only significant for Factor 1 (the general factor), but not in the educational attainment GWAS. The significant genomic loci in the univariate GWASs for the various creativity-related measures and educational attainment are in Supplementary Tables 13 and 14.

### PGS validation

As a verification of the GWASs on the two genetic factors, we tested whether PGSs of the factors could predict creativity phenotypes in the independent STAGE cohort^23–26^. The STAGE cohort comprises approximately 32,000 twin individuals born between 1959 and 1985. In 2019, participants’ saliva samples were collected and genotyped^26^, and non-genotyped monozygotic twins were imputed from their genotyped co-twins if they provided consent. In 2022 and 2023, participants were invited to an online survey including creativity measures. The cohort was not included in any of the GWASs used in our Genomic SEM analyses, and therefore provides a genetically independent sample for testing whether the PGSs generalize to creativity phenotypes.

### PGS calculation

For SNPs on HapMap3 in the GWASs of creativity factors, we used SBLUP^70^ in GCTA^71^ to adjust the estimated SNP effects for LD. We ran SBLUP on the LD matrices, which were calculated from 11,064 unrelated samples randomly selected from the UK Biobank^72^. Using the SBLUP-adjusted SNP effects, we calculated the PGSs for the creativity factors for participants with PLINK2^73^.

### Creativity-related Measures

For PGS predictions, we selected nine creativity(-related) phenotypes in the STAGE cohort: divergent thinking, artistic and scientific achievement, occupational creativity, artistically and non-artistically creative employment, and three openness subscales (aesthetic sensitivity, intellectual curiosity, and creative imagination).

#### Divergent thinking

Participants’ divergent thinking was measured with the Divergent Association Task^3^, which asked participants to list ten nouns that should be as semantically different from each other as possible. The average semantic distance across the words was used to quantify the participant’s divergent thinking ability^74^.

#### Artistic and scientific achievement

Participants’ achievement in creative fields was measured by an adapted Swedish version^75^ of the Creative Achievement Questionnaire^2^. In this questionnaire, participants were instructed to rate their highest achievement (0 = the lowest level, 6 = the highest level) in five artistic fields (music, writing, dance, theater, and visual arts) and two scientific fields (science and invention). By using the maximum scores (ranging from 0 to 6) in the corresponding fields, we calculated participants’ artistic and scientific achievement.

#### Occupational creativity, artistically and non-artistically creative occupations

Following the GWAS on occupational creativity by Kim and colleagues^16^, we derived three creativity-realted phenotypes based on the participants’ occupations. The machine-learning-based continuous variable of occupational creativity ranges between 0 and 1, with a high score indicating that the occupation is more likely to be classified as creative (by the algorithm), and thus, more creative. The other two binary variables identify occupations as artistically or non-artistically creative.Specifically, participants’ occupations were translated from Swedish to English and coded according to the UK SOC 2000 system (with the software Cascot^76^). Values of the occupation-based phenotypes were then assigned to the occupation codes (further details can be found in a previous study by Xia et al.^5^) and participants with the corresponding occupations. For participants reporting multiple occupations, their occupational creativity was the average of the occupational creativity of all listed occupations, and they were regarded as artistically (or non-artistically) creative when any of the listed occupations were artistically (or non-artistically) creative. However, we set the three phenotypes as missing for participants giving more than six occupations.

#### Openness subscales

With the Swedish version of the 30-item Big Five Personality Inventory^77^, we measured three aspects of openness: *aesthetic sensitivity* (e.g., “I am someone who has few artistic interests”), *intellectual curiosity* (e.g., “I am someone who is complex, a deep thinker”), and *creative imagination* (e.g., “I am someone who has little creativity”). Each openness subscale was measured by two items, and each item asked how strongly participants agreed with the corresponding description (0 = completely disagree, 4 = completely agree). We calculated the subscale scores by summing the scores of items in the corresponding subscales (ranging from 0 to 8), after reversing the scores of the false-keyed items (with a lower score indicating higher openness).

### PGS regressions

To examine if PGSs of the two factors could predict creativity-related phenotypes, we fit linear (for continuous phenotypes) and logistic (for binary phenotypes) regressions with covariates of age, sex, and the first 10 principal components (PCs) derived in the STAGE cohort to generate ancestry covariates. As the STAGE cohort is a twin sample, we implemented the sandwich estimators to account for family clustering with the R package “sandwich”. Before regressions, we winsorized outliers (outside the range of *M* ± 3 *SD*) in the continuous creativity-related phenotypes. Except for PCs, all continuous variables in regressions were standardized. The false-discovery rate (FDR) correction was applied to the *P* values to control the FDR at 0.05 across the 18 two-sided tests.

### Genetic correlations

To quantify the genetic associations between creativity and other phenotypes, we employed bivariate LD score regressions^27,29^ and calculated genetic correlations between creativity factors/measures and psychiatric disorders, intelligence, or educational attainment. Similar to the GWASs on creativity-related measures, GWASs on other phenotypes (reported in Supplementary Table 7) were quality controlled, munged with LDSC^29^, and restricted to SNPs on HapMap3^67^. The FDR adjustment was conducted for the 104 two-sided tests on the reported genetic correlations.

### Biological annotation

To further understand the biological pathways connecting the associated SNPs and creativity factors, we conducted gene-based, gene-set, tissue expression, and cell-type expression^78^ analyses with MAGMA^31^ embedded in FUMA^32^. We set the gene annotation window as 10 kb (i.e., SNPs within 10kb of a gene will be assigned to the gene) and excluded the MHC region before running MAGMA. As the default setting of FUMA, we used a Bonferroni correction to account for multiple testing in the biological annotation.

For each latent factor, a gene-based analysis first synthesized the GWAS summary statistics of the SNPs attached to the same gene. A gene-based analysis explored which of the 18,832 genes were significantly associated with the latent factor at the alpha level of 0.05/18,832. Then, a competitive gene-set analysis tested whether the genes in a gene-set were more strongly associated with the phenotype than other genes^31^. For the gene-set analysis on each creativity factor, we applied a Bonferroni-corrected alpha level of 0.05/17,005, as 17,005 gene-sets were tested (including gene-sets curated by FUMA and from MSigDB v2023.1Hs^79^).

FUMA implements the tissue expression analysis with the gene-property analyses^80^ in MAGMA, which examines if genes with higher expressions in one tissue are more strongly associated with the phenotype of interest (i.e., one-sided tests). Our tissue expression analyses were based on the gene expression profiles in 54 tissues from the Genotype-Tissue Expression (GTEx v8) dataset^33^. For each latent factor, a Bonferroni correction was applied to the 54 tests.

Similar to the tissue-expression analysis, the cell-type expression analysis in FUMA uses MAGMA to conduct a gene-property analysis to test if genes with higher expressions in one cell-type are more strongly associated with the phenotype of interest, when the average expression across cell-types in the same region is controlled for. We investigated the gene expressions in 13 types of cells across 105 regions in the brain and spinal cord^34^, which resulted in 1,135 tests (not all regions have all 13 cell types) for each creativity factor. The Bonferroni correction set the alpha level at 0.05/1,135 for each creativity factor.

## Data availability

The summary statistics of existing GWASs used for Genomic SEM and genetic correlation calculation are publicly available or accessible upon request (see Supplementary Tables 1 and 7). Files of LD scores and the reference panel for GenomicSEM are available at https://github.com/GenomicSEM/GenomicSEM/wiki. Upon publication, summary statistics of multivariate GWASs on creativity factors from the current study will be published on GWAS Catalog (https://www.ebi.ac.uk/gwas/), and FUMA results will be accessible at https://fuma.ctglab.nl/browse.

## Code availability

The current study used the R package “GenomicSEM” (v.0.0.5; https://github.com/GenomicSEM/GenomicSEM) in R 4.4.1 for running genome-wide models and multivariate GWASs. Univariate and bivariate linkage disequilibrium score regressions were conducted with LDSC (v.1.0.1; https://github.com/bulik/LDSC). Polygenic score calculation was implemented with SBLUP in GCTA (v.1.94.1; https://yanglab.westlake.edu.cn/software/gcta/#SBLUP) and PLINK2 (v.2.00a6LM, https://www.cog-genomics.org/plink/2.0/). Sandwich estimators were implemented with the R package “sandwich” (v.3.1-1; https://sandwich.r-forge.r-project.org/). FUMA is a publicly online platform (v.2.1.6; https://fuma.ctglab.nl/). Figures were plotted in Python 3.12.11 (https://www.python.org/) with packages of “matplotlib” (v.3.8.4; https://matplotlib.org/), “qmplot” (v.0.3.3; https://github.com/ShujiaHuang/qmplot), “seaborn” (v.0.13.2; https://seaborn.pydata.org/), and “pycirclize” (v.1.10.1; https://github.com/moshi4/pyCirclize). Code for statistical analyses in this study will be available on OSF (https://osf.io/vp7h9/overview?view_only=8af3a97190d34cfdafe5fc0d1a7b0dd2) upon publication.

## Acknowledgement

We thank the Swedish twins for their participation and the Swedish Twin Registry for the data. The present work was supported by the Bank of Sweden Tercentenary Foundation (M11-0451:1) and the Marcus and Amalia Wallenberg Foundation (MAW 2018.0017). The Swedish Twin Registry is managed by Karolinska Institutet and receives funding through the Swedish Research Council under the grant no. 2017-00641. The computations were enabled by resources provided by the National Academic Infrastructure for Supercomputing in Sweden (NAISS) at Uppsala Multidisciplinary Center for Advanced Computational Science (UPPMAX) partially funded by the Swedish Research Council under the grant no. 2022-06725. We thank Sophie Breunig, who kindly helped us with running the genomic structural equation modeling.

